# Quality control of blood plasma for mass-spectrometry proteomic research

**DOI:** 10.64898/2026.01.15.699696

**Authors:** Anastasia V. Vakaryuk, Ivan O. Butenko, Natalia A. Kitsilovskaya, Alexey V. Kovalenko, Veronica D. Gremyacheva, Grigory L. Kozhemyakin, Angelina A. Lebedeva, Nikita M. Baraboshkin, Oleg V. Fedorov, Vadim M. Govorun

**Affiliations:** Research Institute for Systems Biology and Medicine (RISBM)

**Keywords:** mass spectrometry, plasma proteomics, sample quality

## Abstract

Mishandling of samples during blood plasma preparation may lead to biases in proteomic analysis and false interpretation of data. Previous plasma proteome profiling study (*Geyer, Mann et al*., 2019) described proteins that may serve to indicate poor plasma sample quality. We have established a targeted quantitative method for three prominent panels of markers proposed by previous study for exclusion of hemolyzed and coagulation samples. We conducted a model experiment similar to *Geyer, Mann et al*., with both panoramic and targeted detection and investigated the feasibility of application internal control for hemolysis in targeted proteomic analysis by using incorrectly collected plasma. We propose to include plasma sample hemolysis assessment into standard MRM analysis and associated sample exclusion criteria.

## Introduction

Blood plasma proteomics based on mass spectrometric analysis allows for a qualitative and quantitative measurement of protein content in the complex mixture. In addition to the plasma proteins themselves, tissue proteins are secreted into the plasma proteome, changes in the concentrations of which may indicate many diseases. To use a protein biomarker for identification of disease, it is necessary to know precisely that changes in its concentration are due to the patient’s physiological state, and not to errors in the preanalytical stage of sample preparation. The most frequent reasons for sample damage - hemolysis and coagulation. There are many factors that can cause hemolysis or coagulation of blood during sample collection and preparation. In clinics serum is commonly used in diagnostic tests, such as serological testing, discovery of specific proteins and hormones. The widespread use of serum in the clinic is also due to the fact that it is easier to collect and store. Coagulation and subsequent fibrinolysis activate a cascade of biochemical reactions that affect the content of total proteins and analytes significant in clinical diagnostics. Plasma offers a richer source of proteome components and is used to carry out proteomics and metabolomics. The choice of plasma excludes modifications of some analytes due to the coagulation and related with its process, furthermore, plasma volume 15%–20% more compared to serum from the same blood (2).

In clinical laboratories, 40-70% of all spoiled samples are hemolyzed samples (3). Intracellular substances released during red blood cells (RBC) breakdown change the plasma proteomic composition, which can lead to incorrect sample interpretation. Hemolysis interference for such routine biochemistry parameters as: lactate dehydrogenase and aspartate aminotransferase (highly sensitive to hemolysis, free hemoglobin < 0.5 g/L), potassium and total bilirubin (hemoglobin > 1 g/L), alanine aminotransferase, cholesterol, gamma-glutamyltransferase, and inorganic phosphate (changes in concentrations at high levels of hemolysis, hemoglobin: 2.5-4.5 g/L) (4). Therefore, it is common practice in clinical practice to discard spoiled samples based on the hemolysis index (HI) or free hemoglobin concentration (5).

To prevent appointment of false markers, a study on proteome profiling (*Geyer, Mann et al*., 2019) was conducted. In this study three main causes of plasma proteome distortions were considered: coagulation, contamination with platelets or erythrocytes, and, based on the results of a panoramic analysis of samples contaminated in a model experiment, groups of proteins were proposed whose increased or decreased representation may signal these errors in the preanalytical stage of plasma preparation. In this paper, we developed a method for targeted quantification of these proteins and applied it to samples obtained in a similar model experiment, as well as to plasma samples collected under controlled conditions without compliance with the plasma collection procedure.

## Materials and Methods

### Peptide standards selection and synthesis

Based on *Geyer, Mann et al*. results, 17 proteins were selected for targeted quantitation method development (Table 1).

**Table 1.** Proteins for target quantification.

| UniProt ID | Protein Name | Issue | Compared to normal plasma |
| --- | --- | --- | --- |
| P00734 | Prothrombin | Coagulation (or serum) | Downregulated |
| P02656 | Apolipoprotein C-III | Coagulation (or serum) | Downregulated |
| P02671 | Fibrinogen alpha chain | Coagulation (or serum) | Downregulated |
| P02675 | Fibrinogen beta chain | Coagulation (or serum) | Downregulated |
| P02679 | Fibrinogen gamma chain | Coagulation (or serum) | Downregulated |
| P02775 | Platelet basic protein (PBP) | Coagulation (or serum) | Upregulated |
| P07359 | Platelet glycoprotein Ib alpha chain (GP-Ib alpha) | Coagulation (or serum) | Upregulated |
| P07996 | Thrombospondin-1 | Coagulation (or serum) | Upregulated |
| P10909 | Clusterin | Coagulation (or serum) | Upregulated |
| P40197 | Platelet glycoprotein V (GPV) | Coagulation (or serum) | Upregulated |
| P00915 | Carbonic anhydrase 1 | Hemolysis | Upregulated |
| P02042 | Hemoglobin subunit delta | Hemolysis | Upregulated |
| P68871 | Hemoglobin subunit beta | Hemolysis | Upregulated |
| P69905 | Hemoglobin subunit alpha | Hemolysis | Upregulated |
| P21333 | Filamin-A (FLN-A) | Platelet contamination | Upregulated |
| P35579 | Myosin-9 | Platelet contamination | Upregulated |
| Q9Y490 | Talin-1 | Platelet contamination | Upregulated |

For selected proteins a total of 58 candidate tryptic peptide sequences (1-5 per protein) were selected for synthesis and detection test based on their amino acid composition, sequence, position in protein, gene specificity and observed modifications. Peptides were synthesized in pairs of labelled and unlabelled standards. For each peptide a targeted method was developed including retention time, precursor ion, 3-5 fragment ions, declustering potential and collision energy. Unlabelled peptides were used to produce calibration samples of different concentrations (Supplementary 1, 2). Isotopically labelled peptides were added to tryptic digests and calibration samples as internal standards.

### Plasma preparation

To ensure the suitability of the selected markers, we reproduced the experiments as described in methods in *Geyer, Mann et al*. We collected blood from healthy volunteers in VACUETTE K3E K3EDTA tubes. First centrifugation was at 200 g for 10 min, supernatant was harvested for pure plasma extraction. Sediment was centrifugated at 2000 g for 15 min, top layer was discarded. To the sediment were added 4 ml of PBS (1:4) containing 1.6 mg/ml EDTA and centrifuged at 2000 g for 15 min, the supernatant was discarded along with 500 μl of the upper layer of erythrocytes, these steps was repeated twice for extraction pure erythrocytes fraction. Supernatant from the first centrifugation was centrifugated twice at 200 g for 10 min, after the first step were harvested platelet-rich plasma. Supernatant was centrifugated at 2000 g for 15 min for collection of pure plasma. To quantify the degree of contamination with red blood cells, nine dilutions of erythrocytes in pure plasma were made (1:1, 1:4, 10^−1^, 10^−2^, 10^−3^, 10^−4^, 10^−5^, 10^−6^, 10^−7^) (Figure 1, B). In contrast to the original experiment we didn’t use depletion and fractionation.

**Figure 1.**
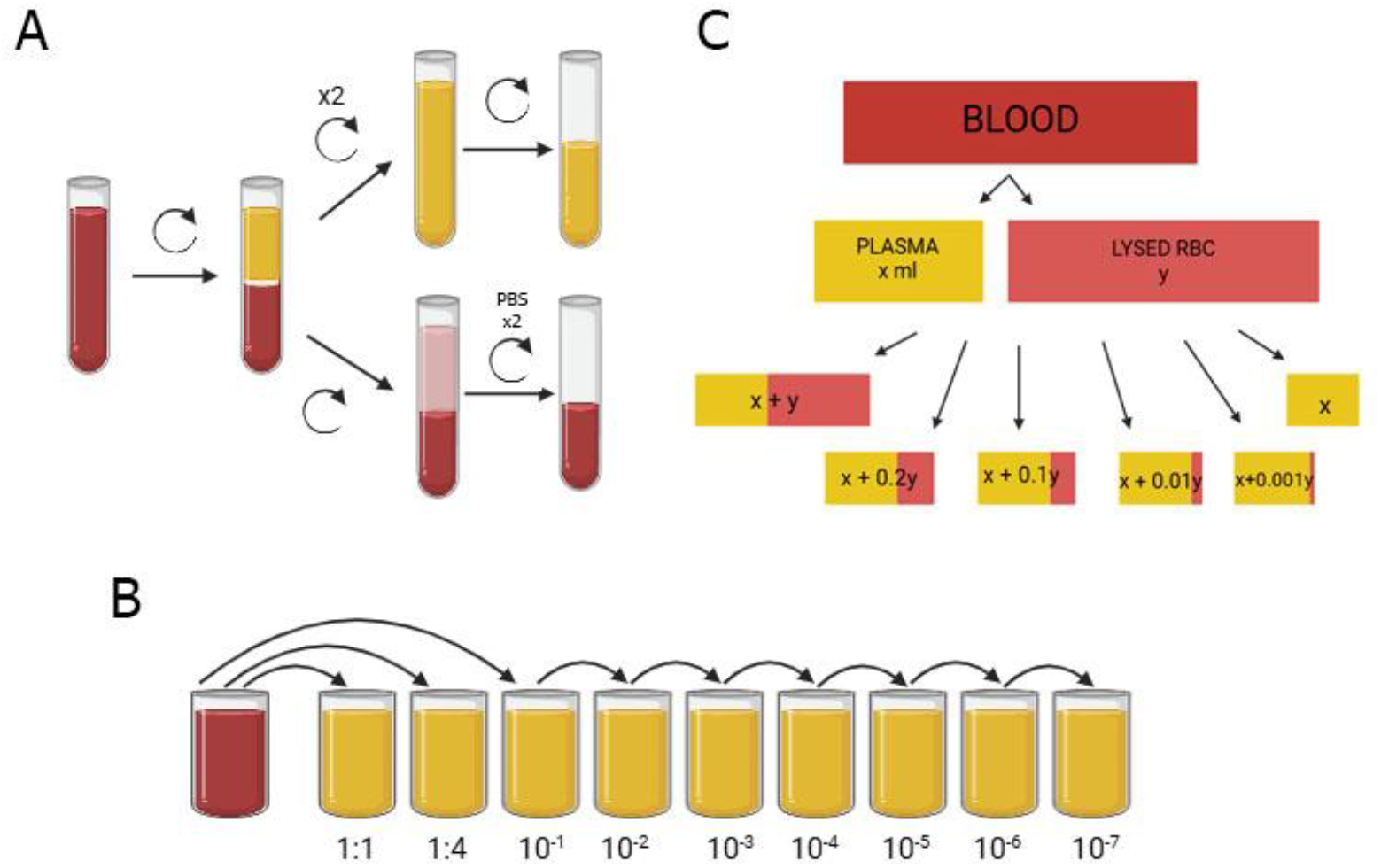
A: obtaining pure RBC and plasma. B: dilutions of erythrocytes in plasma according to the Geyer et al paper scheme. C: dilution of lysed RBC in plasma to simulate a certain degree of hemolysis.

For the second experiment blood was collected from three conditionally healthy donors in VACUETTE K3E K3EDTA 9 ml tubes. Whole blood was centrifuged at 800 *g* for 15 min on 25C in a bucket-rotor to separate plasma from blood cells. Supernatant was repeatedly centrifuged at 3000 g for 20 min to get pure plasma. Bottom layer retained after the first centrifugation was centrifuged at 2000 *g* for 15 min, top layer containing plasma, buffy coat and few erythrocyte was discarded. The erythrocytes were twice washed with PBS buffer (1:2) and centrifuged at 2000 g for 15 min to get pure erythrocytes. Erythrocytes were lysed by mQ added in ratio 1:1. Lysed erythrocytes were added to the plasma in volume, maintaining the ratio of the total plasma volume (x) to the total volume of lysed erythrocytes (y) obtained from one test tube (x + y, x + 0.2y, x + 0.1y, x + 0.01y, x), for comparison of samples with 100, 20, 10, 1, 0.1% hemolysis (Figure 1, C).

Serum was received by collecting blood in vacuum tubes with coagulation activator and gel. Tubes were incubated at room temperature for 30 minutes and centrifugated at 800 g for 15 min. Supernatant was harvested and repeatedly centrifugate at 2000 g 20 min.

Plasma samples were digested with trypsin, preliminary conducting reduction of disulfide bridges with 9 M urea and 20 mM DL-dithiothreitol (DTT) in 300 mM Tris (pH 8.0) and alkylation of cysteines with 100 mM iodoacetamide (IAA). The reaction mixture was diluted with 100 mM Tris to reduce the urea concentration. Trypsin was added at a 1/50 enzyme/substrate ratio twice with an hour gap, and samples were incubated overnight (ON) at 37°C.

After ON incubation trypsin activity was reduced by adding 6% ACN + 0,1% TFA. Synthetic isotope labeled peptides were added to samples for quantification. Plasma purification after enzymatic digestion was performed on a Copure® solid-phase extraction cartridge SPE C18 100 mg/ml. After that samples were dried in HyperVAC-LITE at 2000 rpm, 45°C and reconstituted in 5% ACN 0.1% FA.

### High-pressure liquid chromatography and mass spectrometry

Samples were diluted in 5% ACN and 0,1% formica acid and measured on Orbitrap Exploris 480 combined with UltiMate 3000 HPLC system. Peptides were separated on column Peaky-C18 and eluted with a three-stage linear gradient of 5-65% mobile phase B (80% ACN + 0,1% FA) 60 minutes. The column was washed with 100% mobile phase B for 2 min and post-analytically equilibrated in 5% phase B for 15 min.

Ions were detected in data-dependent analysis. Mass-spectra were acquired in positive ionization mode with a resolution of 60,000 (at m/z 200–1500) for MS and 15,000 for MS/MS scans. The m/z scanning range for MS/MS spectra was selected automatically.

MRM analysis was performed in QTRAP 6500+ Sciex. Peptides were separated in column Restek Ultra AQ C18 (2,1 × 150 mm, particle diameter 3,0 mkl; Restek) at 30°C using a chromatographic system EXION LC-30AD Sciex at a flow rate of 0.3 ml/min for 13 minutes. Synthetic unlabel peptides preliminary were optimase in positive ion registration mode using standard HPLC method optimase for MRM analysis. Optimized parameters: retention time, charge state of the parent ion and 3-5 transition with the maximum signal and collision energy. To the development of MRM analysis, Skyline software was used.

### Spectrophotometric determination of free hemoglobin

Samples of hemolyzed blood were diluted with PBS on 2, 10, 100 times, each dilution was taken in three repetitions. Measured absorption at 415, 380, 450 nm on FlexA-200HTMicroplateReader. The concentration of free hemoglobin were calculated by formula: fHb (mg/dL) = 83.6 (2 × A_415 nm_ − `A`_380 nm_ − `A`_450 nm_) (6), where A415 - oxyhemoglobin absorption, A380 and A450 using for correction for lipids and bilirubins, which may distort the result.

### Data Analysis

MRM analysis was performed at Skyline software (SCIEX). Method for quantitative MRM assay was preliminarily compiled for synthetic peptides using 13C, 15N internal standards, for each pair 4 transitions were selected for more specific determination of the target peptide sequence. The most abundant transition was used for quantification.

Using calibration curves concentration of native peptides was estimated based on unlabel/label peptide peak area. Calibration curve is plotted on lgC ∼ lg(light/heavy). Peptide concentration (fM/mkl) calculated by formula: 10^(a*lg(light/heavy)+b). (C - synthetic peptide concentration in calibration sample, a, b - coefficients in linear equation, light/heavy - 12C/13C or 14N/15N peak area ratio).

MS raw-files from shotgun analysis were analysed by MaxQuant. Searching peptides were performed against reference human proteome from the UniProt database. Trypsin was chosen as an enzyme for digestion, two missed cleavages were allowed. Label free quantification was done with a minimum ratio count - 2.

## Results

### Search for potential markers

Serum and plasma proteome data were provided from shotgun analysis, peptides that between two objects differed by intensity on more than two standard deviations were selected as potential markers (Figure 2). In comparing between our and *Geyer, Mann et al*. coagulation markers (Table 2) we have 7 full coincidences and differing proteins. Fibrinogen and platelet’s proteins differ significantly for plasma and serum because of coagulation mechanism. For some of proteins we found (Hemoglobin subunit alpha,apolipoprotein), changes in concentration between plasma and serum have been noted in other studies (7).

**Table 2.** Proteins in plasma compared to serum in our data vs *Geyer, Mann et al*. markers. (nc - change less than two standard deviations, up - increased in plasma, down - increased in serum)

| ID | Protein name | Mann data | Our data |
| --- | --- | --- | --- |
| P00488 | Coagulation factor XIII A chain | up | up |
| P02671 | Fibrinogen alpha chain | up | up |
| P02675 | Fibrinogen beta chain | up | up |
| P02679 | Fibrinogen gamma chain | up | up |
| P02775 | Platelet basic protein | down | down |
| P02776 | Platelet factor 4 | down | down |
| P07996 | Thrombospondin-1 | down | down |
| P68871 | Hemoglobin subunit beta | nc | up |
| Q13790 | Apolipoprotein F | nc | down |
| P01008 | Antithrombin-III | up | nc |
| P00734 | Prothrombin | up | nc |
| P02656 | Apolipoprotein C-III | up | nc |
| P06276 | Cholinesterase | down | nc |
| P10909 | Thrombospondin-1 | down | nc |
| P33151 | Clusterin | down | nc |

**Figure 2.**
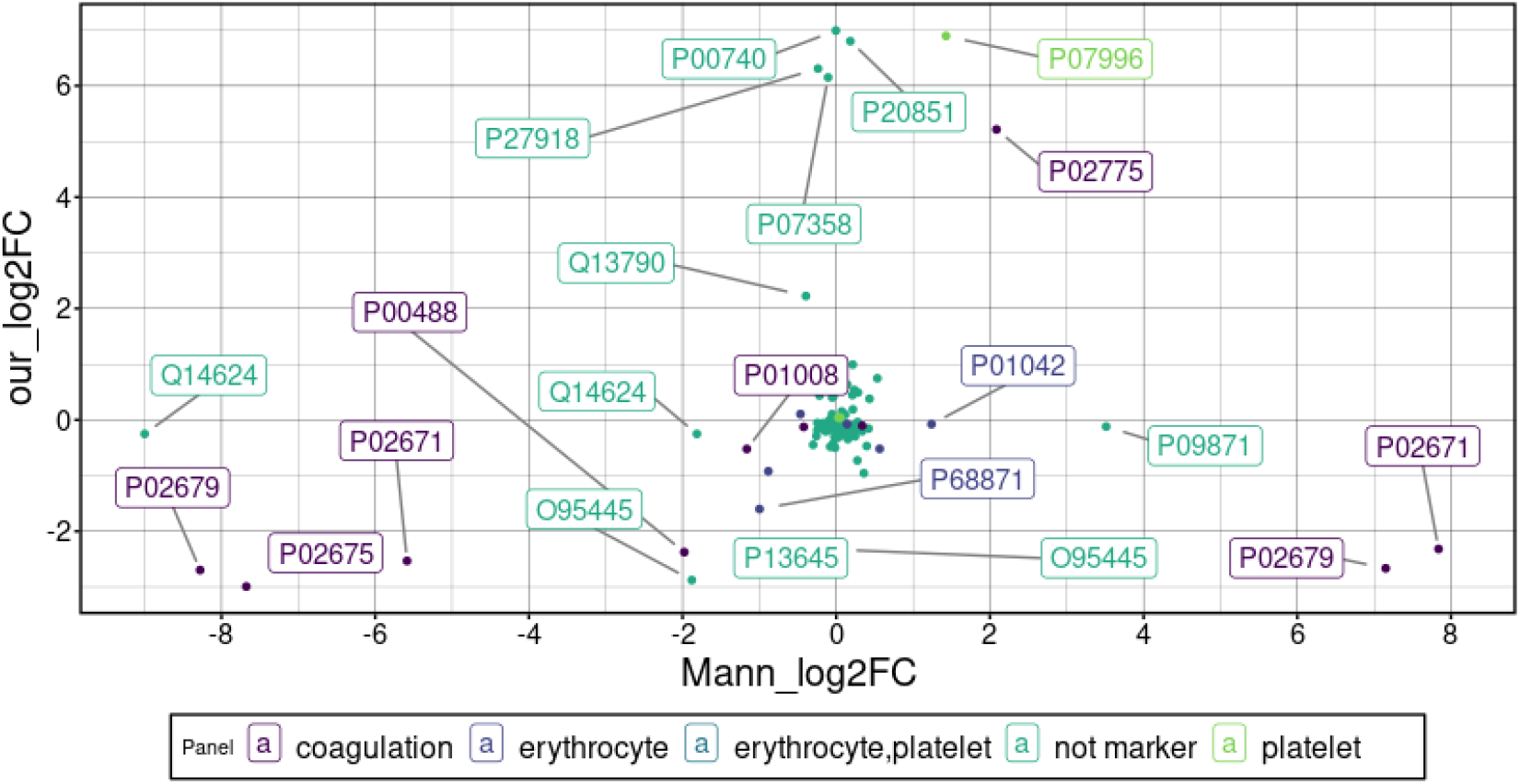
Comparison between serum and plasma proteome in our data and *Geyer, Mann et al*. The colors highlight protein peptides classified as contamination markers in *Geyer, Mann et al*. Infinite values (for proteins found in plasma but not in serum and opposite) were replaced by the extremum on the scale.

During coagulation, fibrinogen is converted into the insoluble protein fibrin, which is removed along with the blood clot during centrifugation. The presence of fibrinogen is considered a key differentiator between plasma and serum, so fibrinogen peptides were chosen as a coagulation marker.

Searching for markers of erythrocyte contamination was carried out in a similar way to that described in the paper: first a panoramic analysis of the erythrocyte proteome was carried out, then 30 high abundance protein with at least a 10-fold higher expression level than in plasma were chosen. 66% of red blood cell (RBC) markers coincided with those proposed in the original paper. (Supplementary 3)

The completeness of proteomes from the objects studied in our study was significantly lower due to absence of depletion and fractionation stage. However, since we choose as potential markers for more abundance proteins in erythrocyte we observe coincidences for some markers regardless of the size of the proteome being studied (Figure 3).

**Figure 3.**
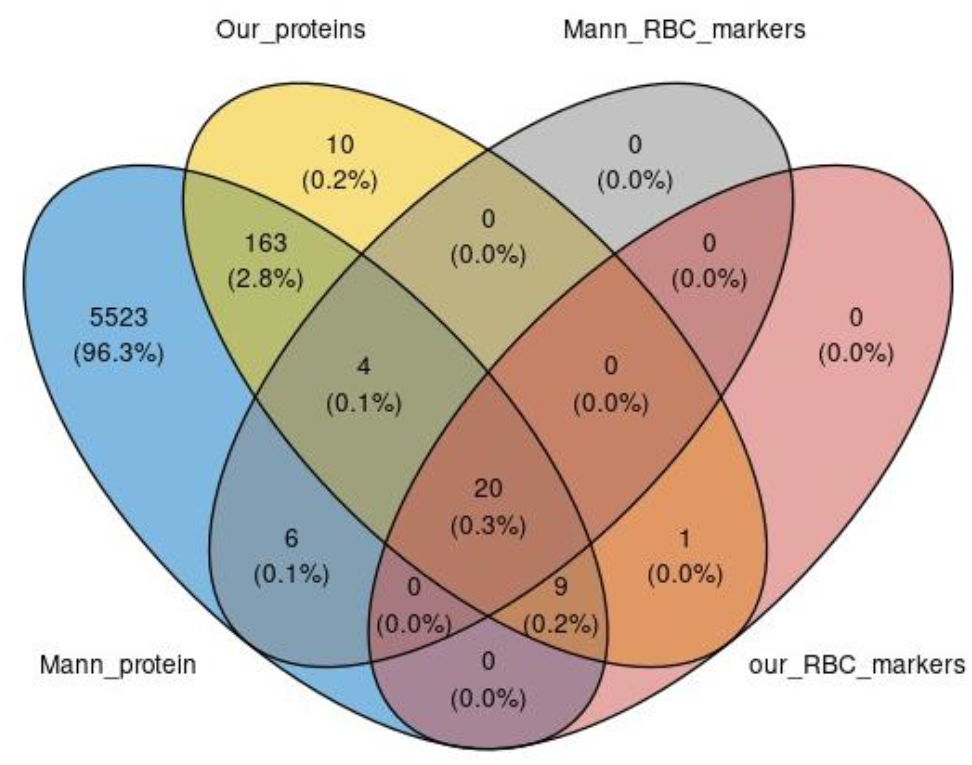
Intersection of sets proteins and markers found in *Geyer, Mann et al*. paper and by our study.

Experiments with RBC diluted in pure plasma imitate different degrees of contamination during hemolysis. A correlation between an increase of the proportion of RBC in plasma and an increase in peptide intensity was observed for hemoglobin subunit peptides, since hemoglobin is the most abundant marker protein for RBC contamination.

### Approval of markers by target analysis

Quantitative proteome analysis performed with synthetic 13C-15N-Arg/Lys labeled peptides added at samples as standards for normalisation. As hemolysis markers were chosen peptides from carbonic anhydrase-1 (2 peptides) and hemoglobin subunit alpha, beta, delta (3 peptides). As a result, one of the carbonic anhydrase peptides and three hemoglobin peptides act as a marker, showing an increase in concentration in samples with an increased degree of hemolysis (Figure 4). Hemoglobin concentration was also measured by spectrophotometry to establish compliance with the quantitative MS method (Figure 5).

**Figure 4.**
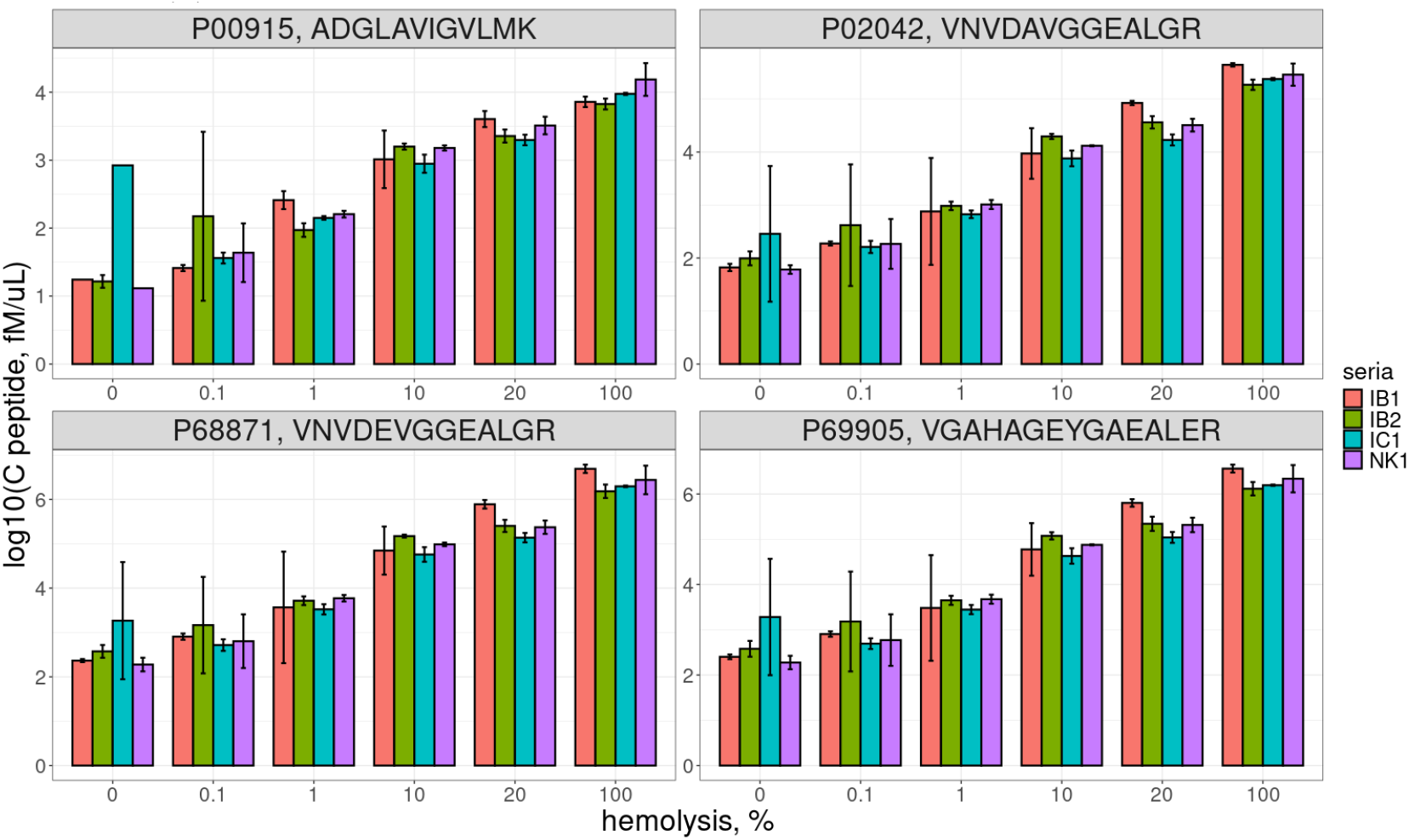
Dependence of marker peptide concentrations on the degree of hemolysis for each donor. A – carbonic anhydrase-1, B, C, D – hemoglobin alpha, delta, and beta. Three technical replicates were performed before the stage of trypsin digestion for each donor.

**Figure 5.**
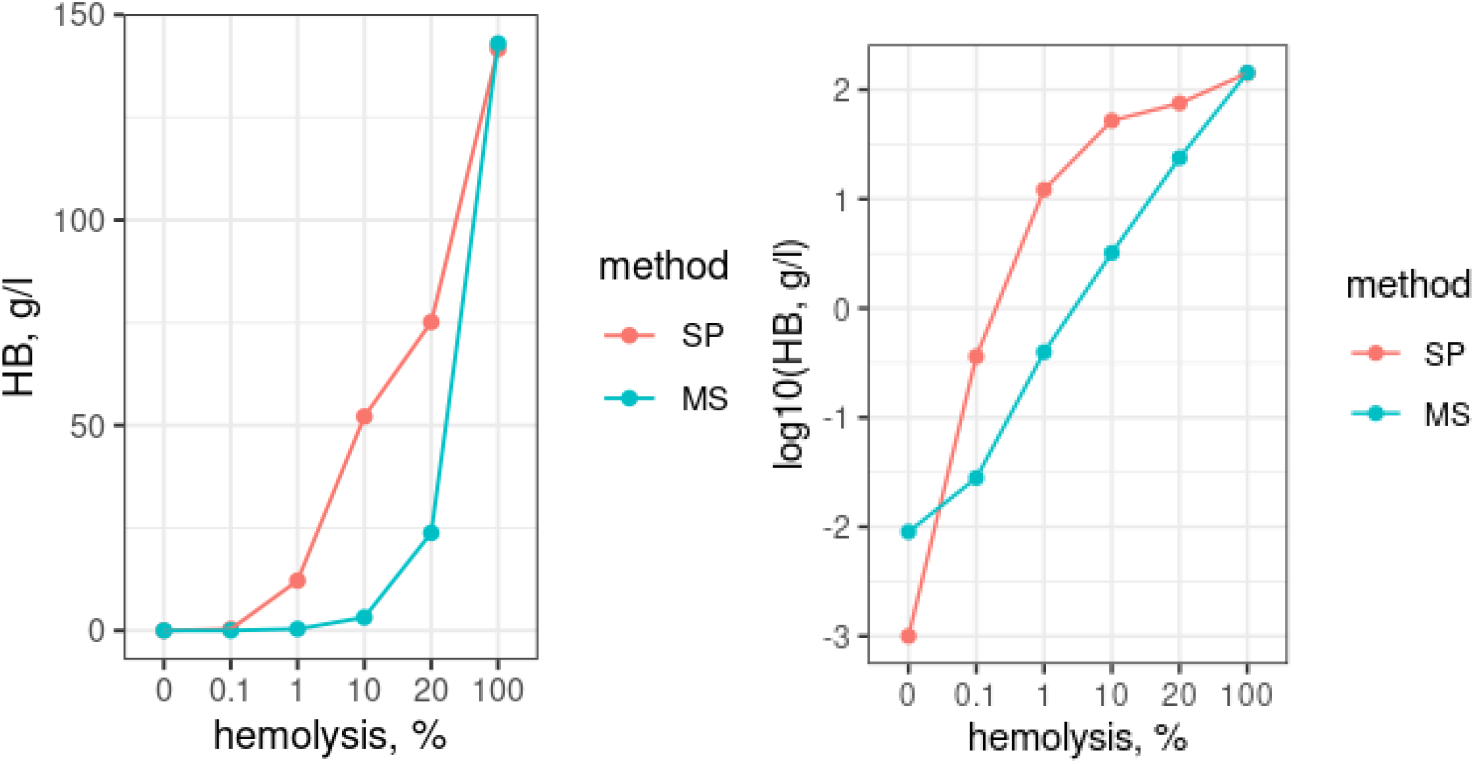
Hemoglobin concentration to hemolysis degree measured by MS and spectrophotometry.

We observed changes in concentration of some clinically significant proteins. From 10% hemolyzed plasma begins a noticeable decrease in protein concentration (Fig. 6). Samples with 10 or more percent of hemolysis are not suitable for proteomics research and should be excluded from analysis.

**Figure 6.**
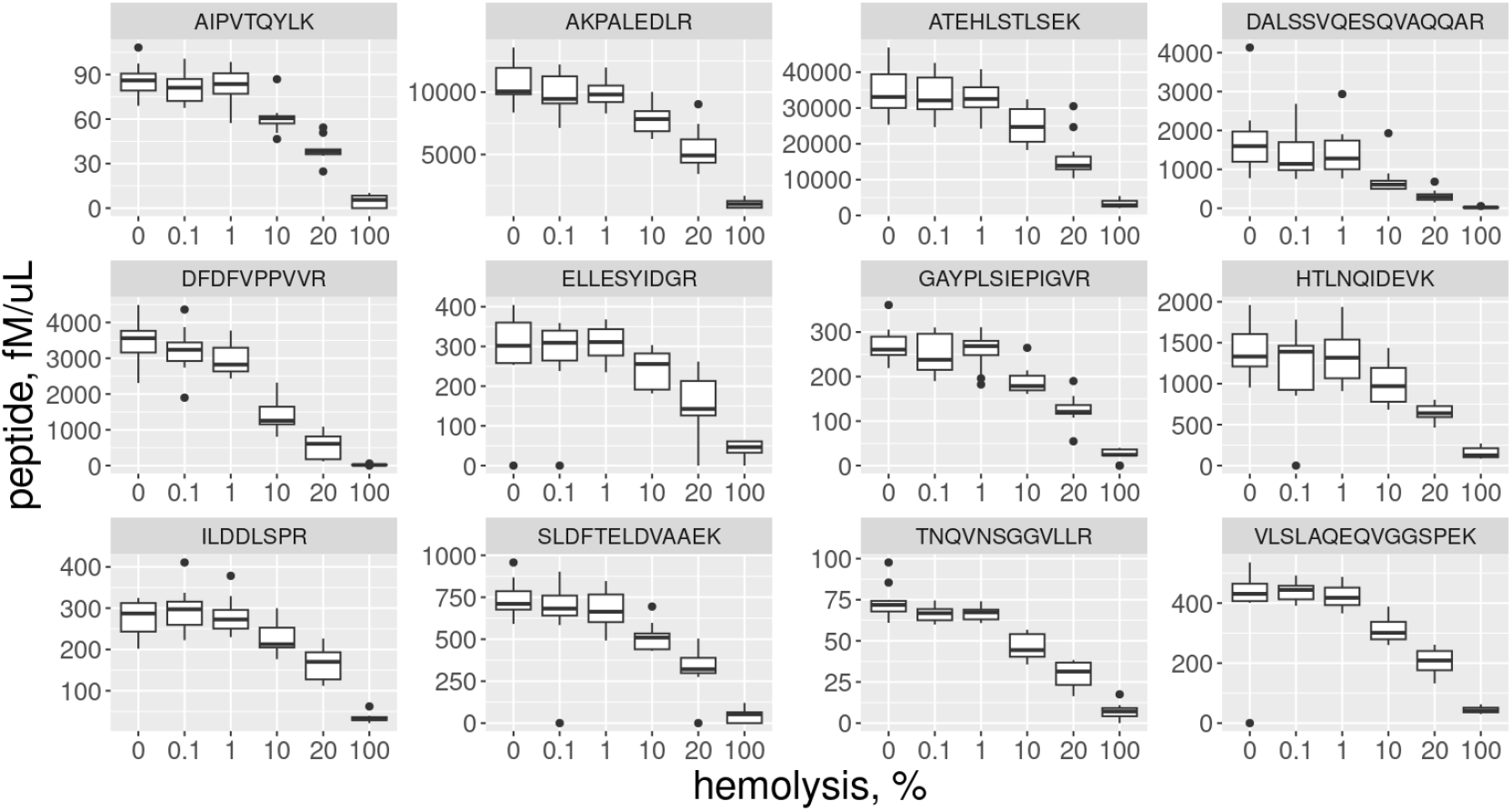
Dependence of the concentration of clinical marker peptides on the degree of hemolysis.

Coagulation affects the proteomic composition of plasma; comparison of the plasma and serum proteome shows differences for some peptides of clinical proteins, for example: Apolipoprotein, Prothrombin, Ceruloplasmin, Plasma protease C1 inhibitor, Afamin, Transthyretin (Figure 7).

**Figure 7.**
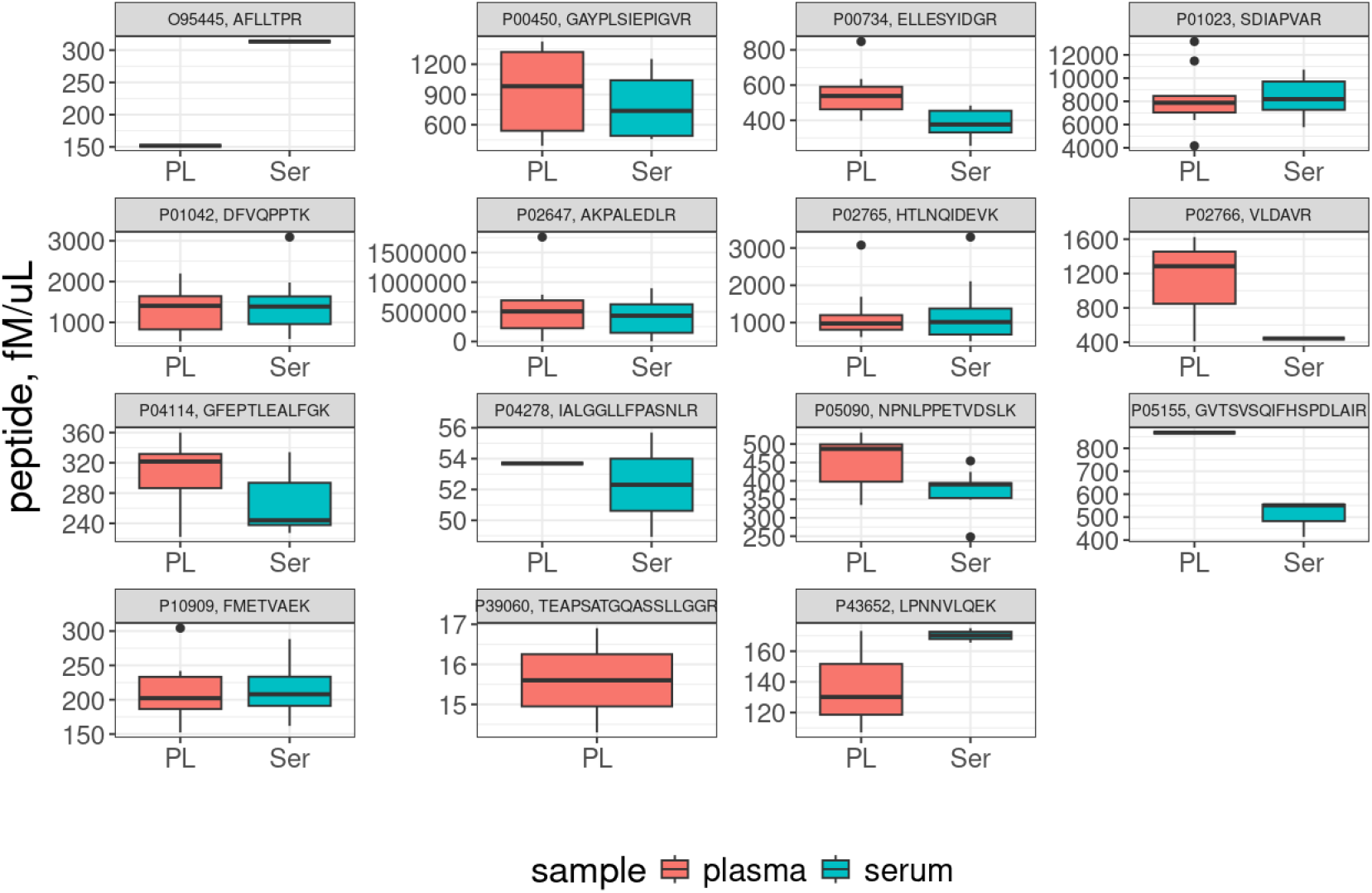
Comparison plasma and serum proteome in MRM.

To test the applicability of the method, a collection of blood plasma received from the hospital was analyzed. Detection of 4 peptides was carried out on the collection, the concentrations of marker peptides were determined by the level of hemolysis of 1% from the previous calculation (Figure 8). We observed strong correlation between hemoglobin’s peptides concentration, less for carbonic anhydrase-1.

**Figure 8.**
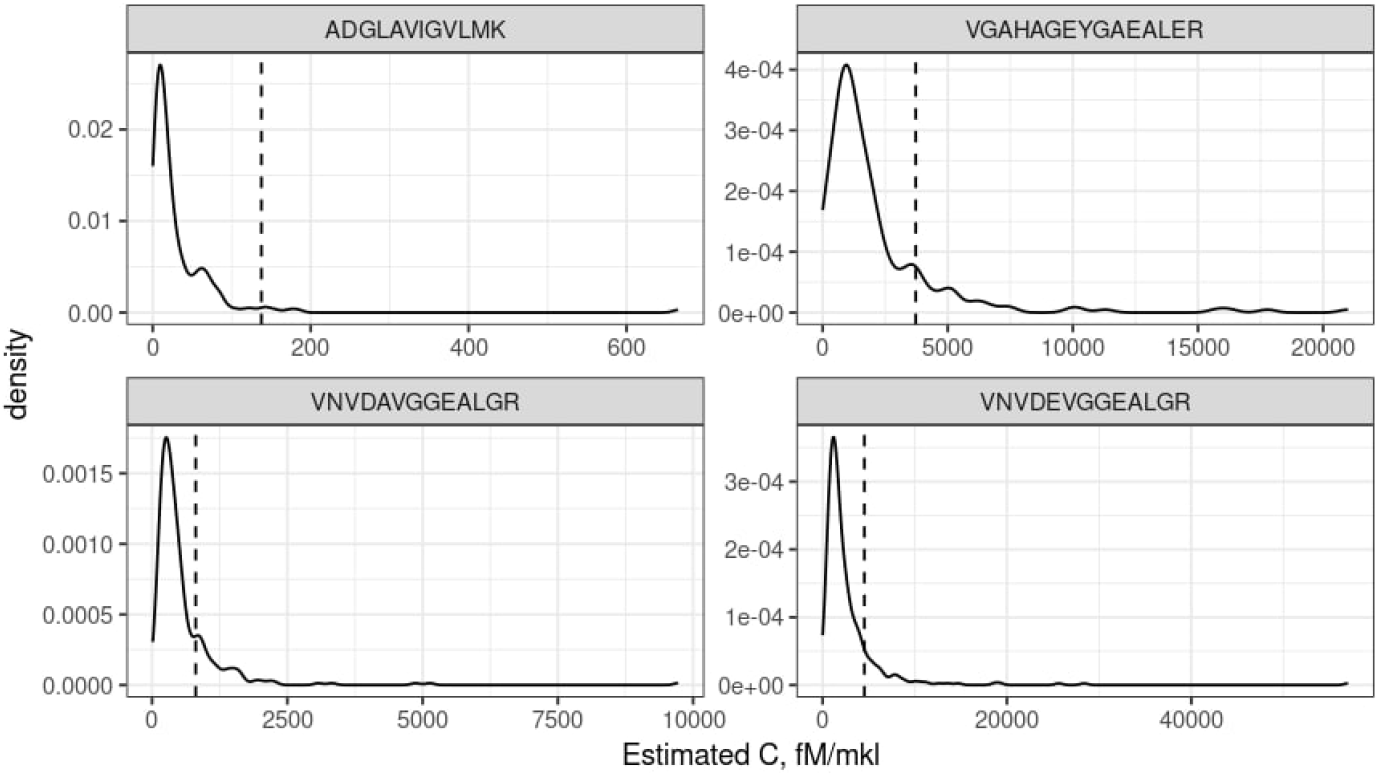
Determination of hemolyzed samples in plasma collection. Dot line - concentration in 1% hemolysis plasma.

## Discussion

Blood plasma is an important biological object for assessing the patient’s condition and health, used in clinical research. Challenges in plasma proteomic studies arise from the large dynamic range of protein concentrations, variability in preanalytical and analytical processes, and biological variability (8). Hemolysis in vitro can cause interference on clinical measurements by changing spectrophotometric values, release of the intracellular components into the sample and sample dilution (9). In clinics conduct their own sample quality tests. To detect hemolysis, the most common occurrence, they determine the free hemoglobin concentration spectrophotometrically. These methods take time and require dedicated equipment. When implementing mass spectrometric proteomics studies for clinical use, analysis time is a crucial factor. Therefore, it is convenient to use quality indicators directly in MS analysis.

To identify contamination markers, we used an approach that is use in the new biomarkers discovery. First, we conducted shotgun analysis to characterize the samples proteomes and selected potential markers among all proteins. Then, we conducted targeted analysis for confirmation. This approach allowed us to select coagulation and hemolysis markers and determine a contamination level due to erythrocytes lysis, which can be used to exclude samples from analysis.

## Funding

The work was funded by Federal Service for Surveillance on Consumer Right Protection and Human Wellbeing grant № 124021600055-6

## Notes

### Competing Interest Statement

The authors have declared no competing interest.

