## Supplemental 3 for "Quality control of blood plasma for mass-spectrometry proteomic research"

|  | **confirmed markers** | | **differents** | |
| --- | --- | --- | --- | --- |
|  | ***Protein ID*** | ***Protein name*** | ***Protein ID*** | ***Protein name*** |
| **erythrocyte contaminations** | P00558-2 | Phosphoglycerate kinase 1 | P02768-1 | Albumin |
|  | P00915 | Carbonic anhydrase 1 | P07477 | Serine protease 1 |
|  | P00918 | Carbonic anhydrase 2 | P16452-3 | Protein 4.2 |
|  | P02042 | Hemoglobin subunit delta | P10599-2 | Thioredoxin |
|  | P02549-2 | Spectrin alpha chain | P11166 | Solute carrier family 2, facilitated glucose transporter member 1 |
|  | P02730-2 | Band 3 anion transport protein | P27105 | Stomatin |
|  | P04040 | Catalase | P37840-2 | Alpha-synuclein |
|  | P04406-2 | Glyceraldehyde-3-phosphate dehydrogenase | P60981 | Destrin |
|  | P07738 | Bisphosphoglycerate mutase | P69892 | Hemoglobin subunit gamma-2 |
|  | P11171-6 | Protein 4.1 |  |  |
|  | P11277-2 | Spectrin beta chain, erythrocytic |  |  |
|  | P16157-14 | Ankyrin-1 |  |  |
|  | P30041 | Peroxiredoxin-6 |  |  |
|  | P30043 | Flavin reductase (NADPH) |  |  |
|  | P32119 | Peroxiredoxin-2 |  |  |
|  | P63261 | Actin, cytoplasmic 1 |  |  |
|  | P68871 | Hemoglobin subunit beta |  |  |
|  | P69905 | Hemoglobin subunit alpha |  |  |

Table 1. Markers of RBC contamination
